# GGSDT: A unified signal detection framework for confidence data analysis

**DOI:** 10.1101/2022.10.28.514329

**Authors:** Kiyofumi Miyoshi, Shin’ya Nishida

## Abstract

Human decision behavior entails a graded awareness of its certainty, known as a feeling of confidence. Until now, considerable interest has been paid to behavioral and computational dissociations of decision and confidence, which has raised an urgent need for measurement frameworks that can quantify the efficiency of confidence rating relative to decision accuracy (metacognitive efficiency). As a unique addition to such frameworks, we have developed a new signal detection theory paradigm utilizing the generalized gaussian distribution (GGSDT). This framework evaluates the observer’s internal standard deviation ratio and metacognitive efficiency through the scale and shape parameters respectively. The shape parameter quantifies the kurtosis of internal distributions and can practically be understood in reference to the proportion of the gaussian ideal observer’s confidence being disrupted with random guessing (metacognitive lapse rate). This interpretation holds largely irrespective of the contaminating effects of decision accuracy or operating characteristic asymmetry. Thus, the GGSDT enables hitherto unexplored research protocols (e.g., direct comparison of yes/no versus forced-choice metacognitive efficiency), expected to find applications in various fields of behavioral science. This paper provides a detailed walkthrough of the GGSDT analysis with an accompanying R package (*ggsdt*).

## Introduction

Humans have introspective awareness of their own decision correctness, and this feeling of confidence performs a key role in guiding future adaptive behavior (e.g., Boldt et al., 2019; Desender et al., 2019; Guggenmos et al., 2016). While successful behavioral guidance rests on the confidence’s predictability of decision correctness (i.e., metacognitive accuracy), studies have accumulated rich evidence indicating their potential dissociations (e.g., Komura et al., 2013; Maniscalco & Lau, 2015; Miyamoto et al., 2017; Rounis et al., 2010; Weiskrantz, 1974). These dissociations have attracted growing attention in recent years since unraveling their possible causes is expected to provide a breakthrough in broad areas of behavioral science (e.g., Hoven et al., 2019; Michel, 2022; Peters, 2022; Rouault et al., 2018; Rouy et al., 2021).

The quantitative characterization of the decision-confidence dissociation requires mathematical frameworks that can evaluate the accuracy of metacognitive monitoring relative to objective decision accuracy (a concept termed metacognitive efficiency). The present paper sets out to propose a general analysis framework to this end upon a solid methodological foundation of signal detection theory (SDT; see Wixted, 2020 for a review). The theory portrays latent evidence distributions that would best describe observed decision-making behavior, the parameters of which effectively characterize the subject’s decision performance (e.g., standardized mean difference, standard deviation [SD] ratio, etc.).

There have been two major branches of detection theory—classic SDT (e.g., Egan & Clarke, 1956; Green & Swets, 1966; Macmillan & Creelman, 2005) and meta-SDT (e.g., Clarke et al., 1959; Galvin et al., 2003; Maniscalco & Lau, 2012)—that are predicated on different theoretical assumptions for different analysis purposes. As outlined below, each of these frameworks has distinct benefits (and limitations) that the other does not share. While the classic SDT enables more flexible modeling implementations, only the meta-SDT serves to evaluate the observer’s metacognitive efficiency. Therefore, in order to incorporate the advantages of these two frameworks, we have developed a new analysis framework termed generalized gaussian signal detection theory (GGSDT), with a complementary analysis package available on the R platform.

The following sections will develop with a picture of basic psychology experiments accompanied with confidence rating (**Figure 1**), the data from which have traditionally been analyzed by the classic SDT or meta-SDT (**Figures 2, 3**; see **Materials and methods** for details). With these as theoretical foundations, we propose our GGSDT framework through simulations and fittings to empirical data. To anticipate, the GGSDT quantifies metacognitive efficiency by a kurtosis parameter, referring to the proportion of the gaussian ideal observer’s confidence disturbed with random lapse. This interpretation is generalizable over different internal variance structures as the GGSDT accommodates itself to asymmetric receiver operating characteristics (ROCs) through a scale parameter. With these properties, the GGSDT would be a valuable measurement tool for broad behavioral science disciplines.

**Figure 1.**
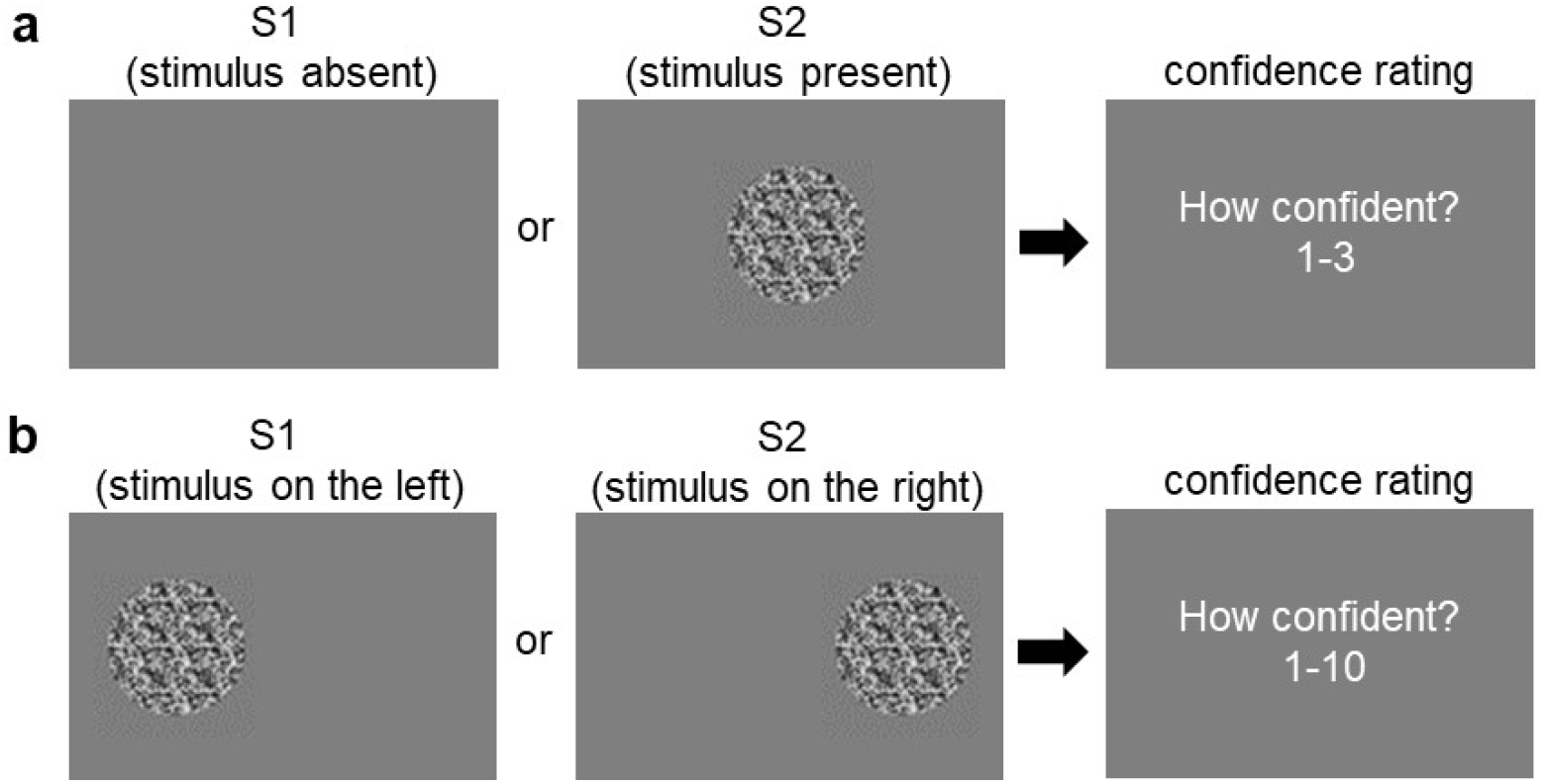
Two different formats of experiments. (**a**) An example of yes/no visual detection. The observer decides if the external world state is S1 (stimulus absent) or S2 (stimulus present), and then gives a confidence rating. Usually, the state of stimulus presence gives greater internal evidence variability than the state of stimulus absence, leading to asymmetric type-1 ROC data (see **Figure 2a**). (**b**) An example of left/right two-alternative forced-choice (2AFC) visual discrimination. The observer discriminates if the external world state is S1 (grating on the left) or S2 (grating on the right), followed by a confidence rating. Because of the symmetric structure of the S1 and S2 states, symmetric type-1 ROC data are usually observed in 2AFC experiments.

**Figure 2.**
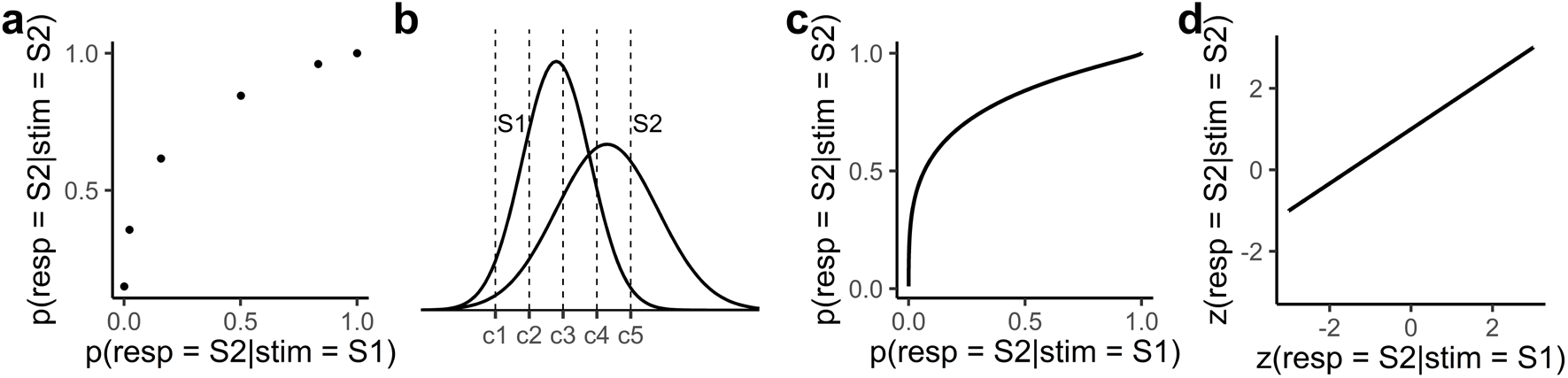
Classic gaussian SDT analysis, standing upon **Figure 1a**. (**a**) Example type-1 ROC data showing 6 pairs of hit and false alarm rates. In counting hits and false alarms, the point more on the left includes trials assigned with greater favor for S2 response (e.g., the left-most point only counts the highest confidence S2 responses). (**b**) Internal evidence under S1 and S2 states is assumed to be distributed from one trial to another. Having a parameter for the S2/S1 SD ratio, the classic SDT allows flexible variance structures. The internal decision space is demarcated by decision criteria (c1-c5) into 6 regions, which correspond to 6 possible response classes (highest confidence S1 to highest confidence S2). (**c**) Type-1 ROC prescribed by the classic gaussian SDT. Because of the unequal variance in the internal decision space, the ROC exhibits asymmetry regarding the major diagonal. (**d**) Type-1 ROC in z-transformed space. The slope of zROC is determined by the SD ratio of the internal distributions.

**Figure 3.**
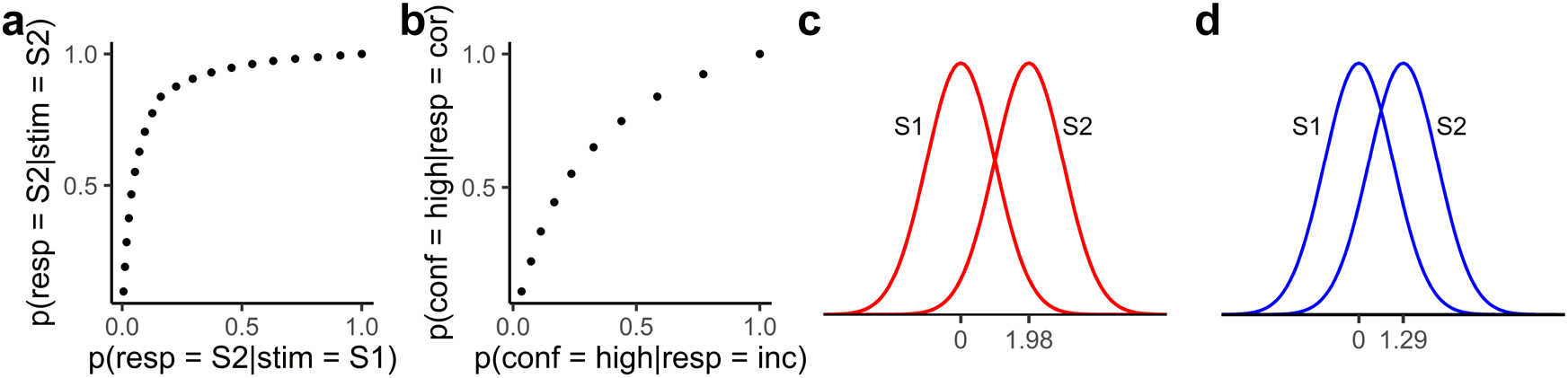
Gaussian meta-SDT analysis, standing upon **Figure 1b**. (**a**) Example type-1 ROC data, showing the discrimination of external world states with both objective decision and confidence rating. (**b**) Example type-2 ROC data, demonstrating the discrimination of the observer’s own decision correctness with confidence rating. (**c**) Objective decision data (a pair of hit and false alarm rates) is fitted by so-called type-1 distributions. The standardized mean difference of these distributions constitutes a parametric measure of objective decision accuracy known as d’. (**d**) Type-2 ROC data are fitted by so-called type-2 distributions. The standardized mean difference of these distributions gives a parametric measure of metacognitive accuracy known as meta-d’. The meta-d’/d’ ratio (m-ratio) is often employed to quantify the observer’s metacognitive performance relative to objective decision accuracy (metacognitive efficiency). Note that the equal variance assumption is necessary in the fitting of type-1 and type-2 distributions and the meta-SDT has difficulty in explaining asymmetric type-1 ROC data.

## Results

### Proposal of generalized gaussian signal detection theory (GGSDT)

Metacognitive performance has traditionally been measured with the analysis of type-2 ROC data. We have found, however, that type-1 ROC data also have a systematic connection to the observer’s metacognitive performance, which inspired us for designing the GGSDT (**Figure 4**). In constructing empirical type-1 ROC, the middle data point (nth point from the left for n-level confidence data) is solely determined by objective decisions (i.e., a pair of hit and false alarm rates), whereas the efficiency of metacognition is reflected in the other data points. If confidence rating is not diagnostic of objective decision correctness at all, type-1 ROC data consist of straight line segments that connect the middle data point (10th point from the left in **Figure 4a**) to (0, 0) and (1, 1). Under a given objective accuracy, data points other than the midpoint expand outward with growing metacognitive efficiency (**Appendix 1**).

**Figure 4.**
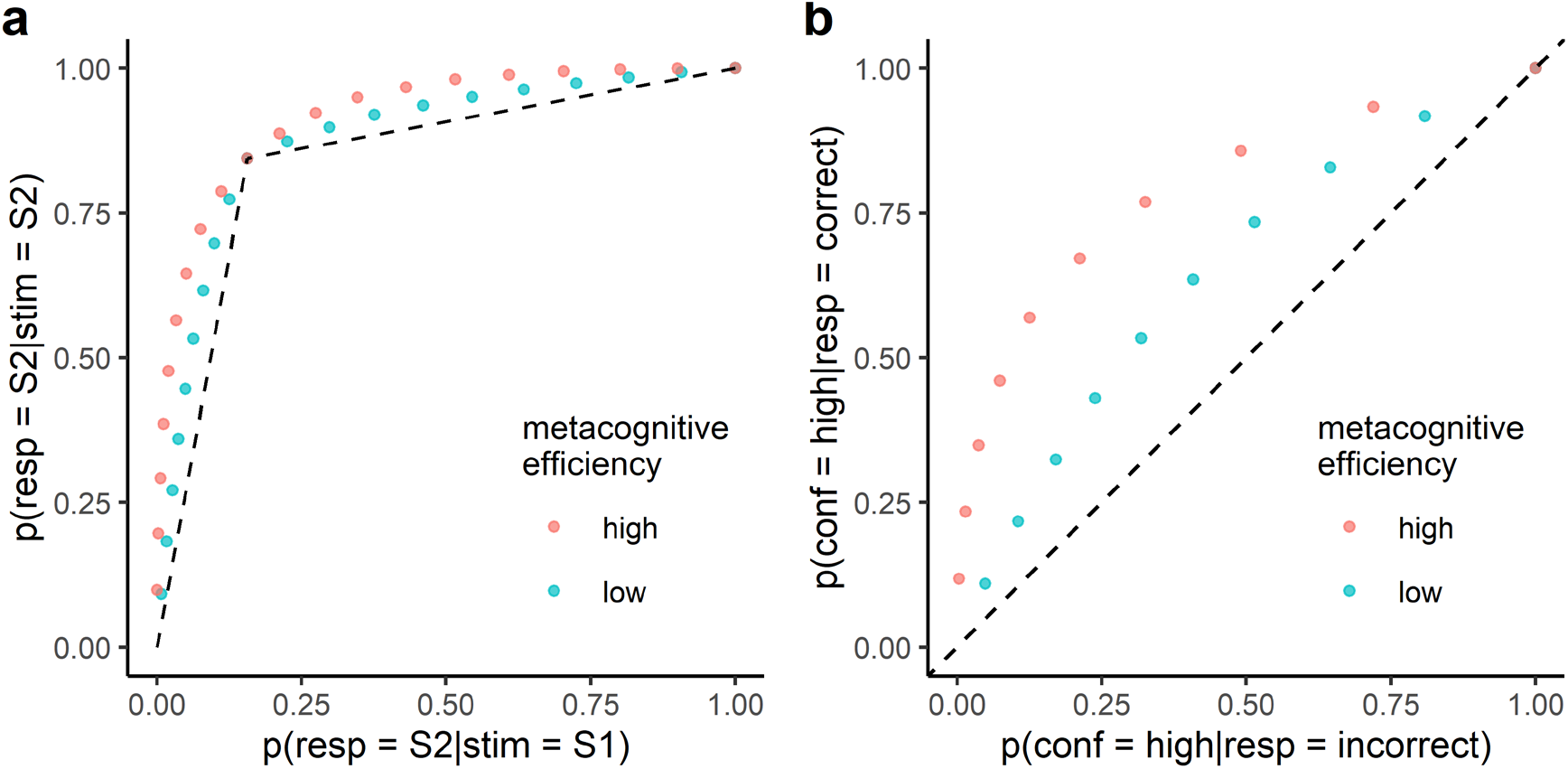
The crosstalk between type-1 versus type-2 operating characteristics illustrated with 10-level confidence rating data. Blue and red example data are equivalent in terms of objective decision accuracy yet are different in metacognitive efficiency. (**a**) In type-1 ROC space, the middle data point (point tenth from the left) solely represents objective decision accuracy, while metacognitive performance is reflected by the expanding curvature of the other data points. (**b**) In type-2 ROC space, metacognitive performance is reflected by the area under the curve. Dashed lines indicate the performance characteristics under zero metacognitive efficiency.

Importantly, from the classic SDT perspective, this expanding curvature is controlled by internal distributions’ kurtosis (Miyoshi et al., 2022). The greater the kurtosis, the smaller the expansion becomes, indicative of the lower type-2 hit rate and the higher type-2 false alarm rate. Therefore, by estimating distribution kurtosis from type-1 ROC data, one can evaluate the observer’s metacognitive performance without referring to type-2 ROC.

The GGSDT is simply a classic SDT with a generalized gaussian distribution assumed for internal distributions (**Figures 5, 6**). The generalized gaussian distribution is defined with three parameters, the mean (μ), scale parameter (α), and shape parameter (β). The followings describe the formulae for calculating the summary statistics of the generalized gaussian distribution, where G() represents the gamma function. Since distribution kurtosis is solely determined by β, hereafter we use it as an index of metacognitive efficiency.^1^

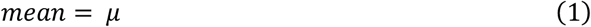

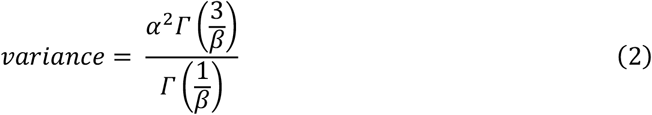

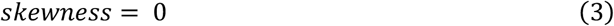

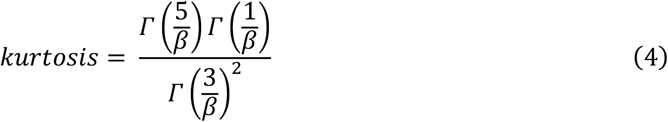

**Figure 5.**
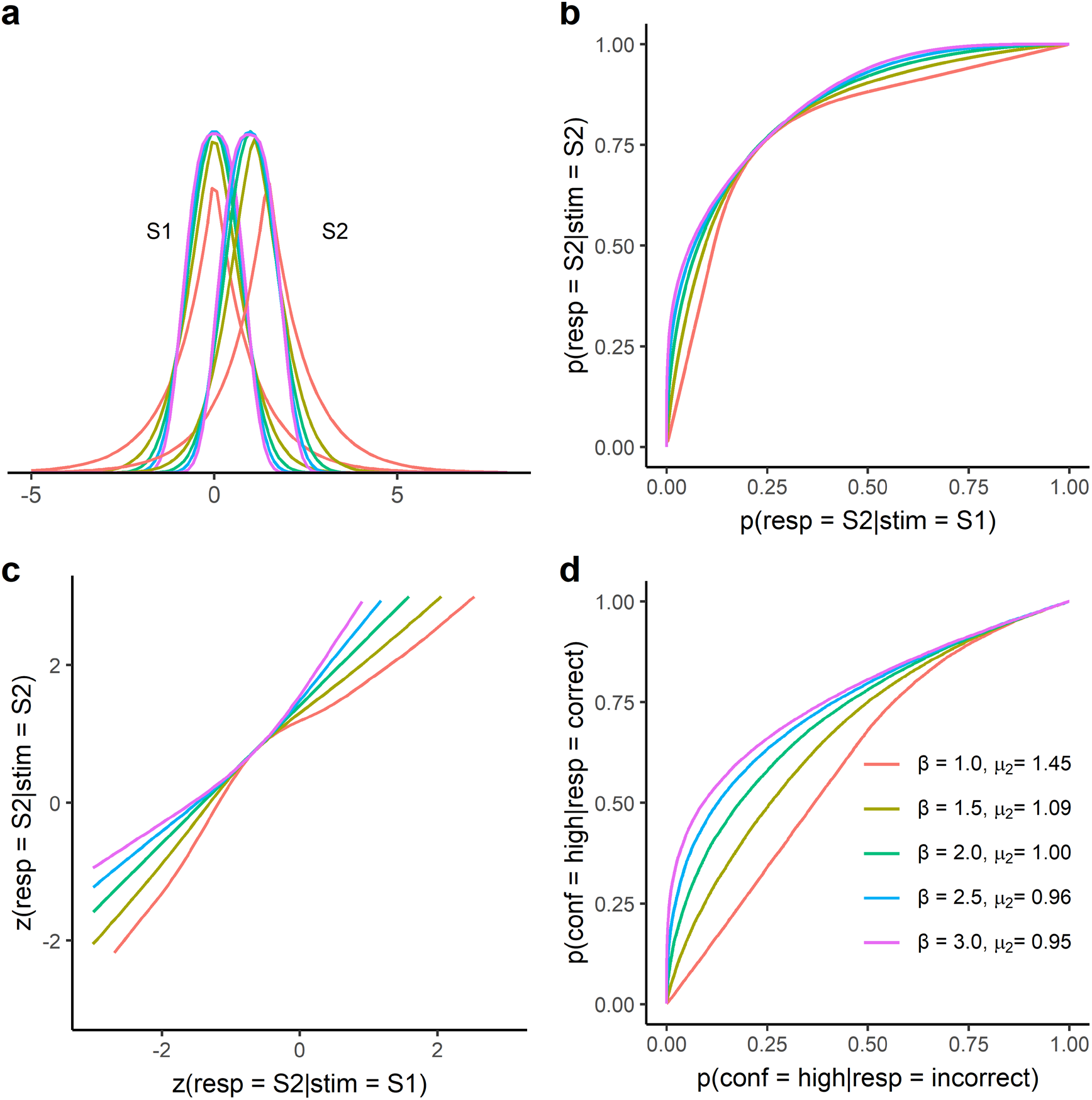
The behavior of the GGSDT under the equal variance scenario (α_2_ is set at 1.0). (**a**) Internal distributions of the GGSDT with different values of β. The μ_2_ parameter is also manipulated to emulate equal objective decision accuracy across the conditions of different β values. (**b**) Type-1 ROCs prescribed by the GGSDT. Different β values lead to changing extent of expanding curvature, reflecting varying levels of metacognitive efficiency. (**c**) Type-1 zROCs prescribed by the GGSDT. β values > 2 give an upright U-shape whereas β values < 2 lead to an inverted U-shape. (**d**) Type-2 ROCs prescribed by the GGSDT. Different β values result in varying sizes of the area under the curve, which signifies changing levels of metacognitive efficiency.

**Figure 6.**
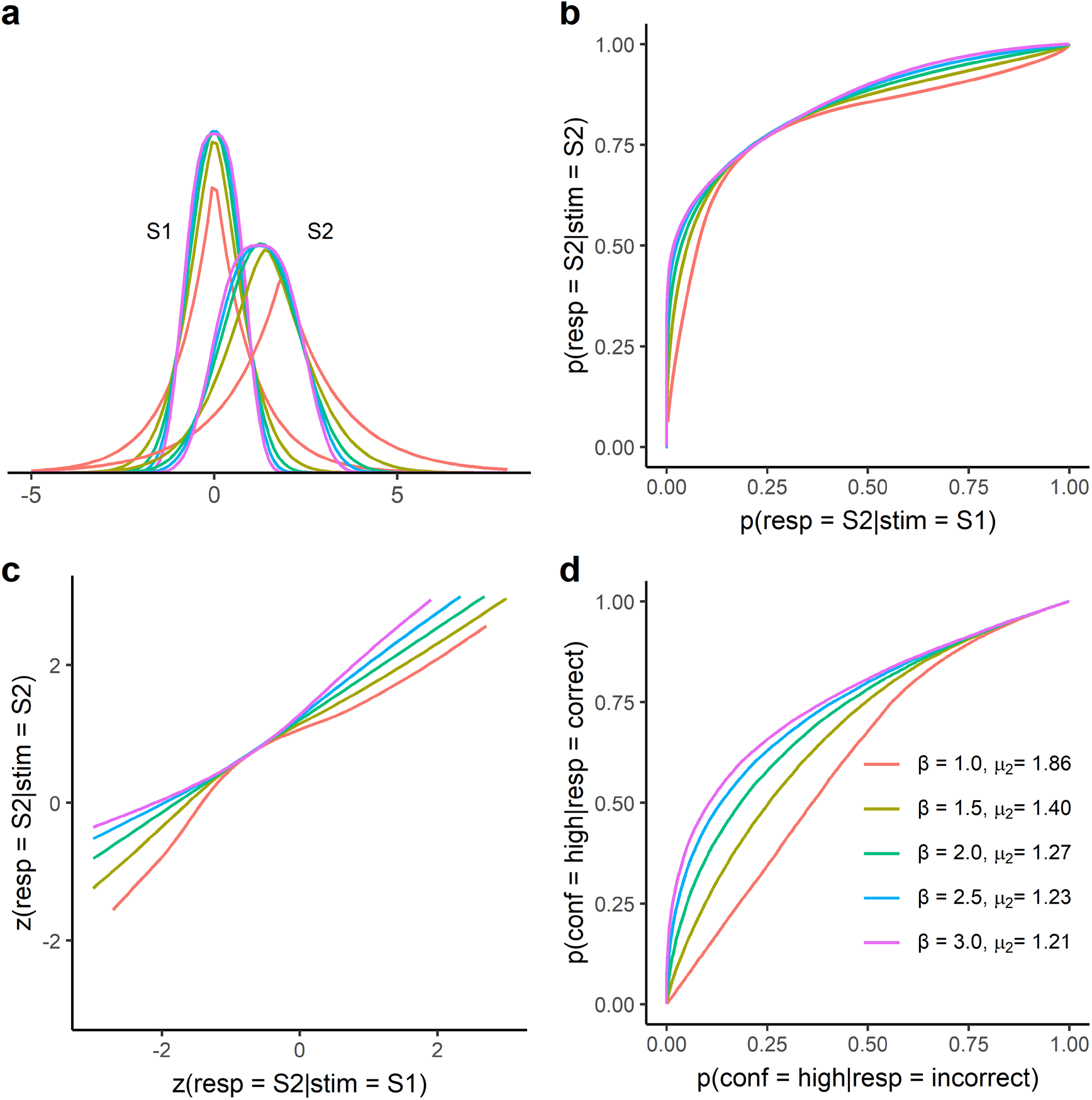
The behavior of the GGSDT under the unequal variance scenario (α_2_ is set at 1.5). (**a**) Internal distributions of the GGSDT with different values of β. (**b**) Type-1 ROCs. (**c**) Type-1 zROCs. (**d**) Type-2 ROCs.

The GGSDT model estimates the mean and the scale parameter of the S2 distribution (μ_2_ and α_2_), while those of the S1 distribution are scaled to be 0 and 1 respectively. The model also estimates the shape parameter β, which is assumed to be identical for S1 and S2 distributions in the current implementation. Due to this formalization, α_2_ represents the SD ratio of two internal distributions (see equation 2). The generalized normal distribution agrees with the Laplace distribution when β = 1, the gaussian distribution when β = 2, and the uniform distribution when β = ∞. Therefore, for example, a β estimate of 2 indicates that the observer is as metacognitively efficient as the classic gaussian SDT. This is conceptually analogous to the gaussian meta-SDT analysis, where the pattern of m-ratio = 1 indicates that the observer’s metacognition is as efficient as the classic equal variance gaussian SDT. Still, the GGSDT allows more flexible model implementations, unrestricted by the equivariance assumption.

### Behavioral characteristics of the GGSDT

**Figure 5** demonstrates the performance characteristics of the GGSDT under varying β values with α_2_ fixed at 1.0. To better visualize the β’s relevance to metacognitive efficiency, we chose to plot the figures emulating constant objective decision accuracy. Specifically, we fixed the GGSDT’s type-1 ROCs regarding their intersection with the major diagonal, at which objective decision accuracy is set at 76% (rendering constant objective decision accuracy under unbiased decision criterion). Since β changes the scale of the decision space (SD of S1 and S2 distributions) through equation 2, we modulated μ_2_ to fixate the intersection point (see **Appendix 2** for model behavior without μ_2_ modulation).

**Figure 5a** shows the GGSDT’s internal distributions, where the kurtosis of the distributions changes depending on the β value. Smaller β (i.e., greater kurtosis) gives smaller expansion in type-1 ROC space (**Figure 5b**) as well as inverted U curvilinearity in type-1 zROC space (**Figure 5c**), which are mirrored in type-2 ROC space as impaired metacognitive efficiency (**Figure 5d**). Importantly, these basic properties are preserved if unequal variance structure is introduced in the internal decision space (**Figure 6**). The GGSDT’s behavior is displayed here for a range of β values with α_2_ fixed at 1.5. As with the equal variance situation above, the type-1 ROCs are plotted so that objective accuracy stays constant at 76% at their intersection with the major diagonal. Notably, these figures collectively demonstrate that, under a certain objective decision accuracy, a given value of β yields nearly constant type-2 operating characteristics regardless of the α_2_ parameter (**Appendix 2** also shows β’s SD-ratio-independent property from a different angle). Note that these visualizations are only to help understand the functions of the GGSDT parameters, and actual model fitting is not done in such a way that the objective decision data point is fixed. Yet, the visual comprehension of the model’s behavioral characteristics provides useful insights in interpreting the GGSDT data analysis presented in the following sections.

It is noteworthy that the inverted U-shaped zROC (seen with β < 2) has been observed in human behavioral experiments, particularly in the domain of visual perception (Miyoshi et al., 2022; Shekhar & Rahnev, 2021). Also, the upright U-shaped zROC (seen with β > 2) has occasionally been found in recognition memory studies (e.g., Hilford et al., 2002; Rotello et al., 2000; Slotnick et al., 2000; Slotnick, & Dodson, 2005; Starns et al., 2013; Yonelinas, 1997; Yonelinas & Parks, 2007). Therefore, the GGSDT prescribes a performance measurement under the constraint that is qualitatively consistent with human behavioral characteristics.

In summary, the GGSDT does not explicitly distinguish between objective decision and confidence rating, describing both with a common set of internal distributions. Yet, it captures the observer’s metacognitive efficiency through the β parameter while accommodating unequal variance structure with the α_2_ parameter. Accordingly, the GGSDT combines the good modeling transparency of the classic SDT with the metacognitive evaluation capability of the meta-SDT.

### Fitting to simulation data

Next, to demonstrate GGSDT’s properties in parameter estimation, we fit the model to the data sampled from the classic equal variance gaussian SDT (d’ = 1.5). First, we have sampled 200,000 observations from S1 and S2 distributions and simulated objective decision and confidence rating according to the following optimal Bayesian rule. Here, p(S1) and p(S2) represent the prior probability of S1 and S2 states (assumed to be 0.5 in all following implementations), p(x|S1) and p(x|S2) are the likelihood of observation x given S1 and S2 states, and p(S1|x) and p(S2|x) are the posterior probability of S1 and S2 states given observation x.

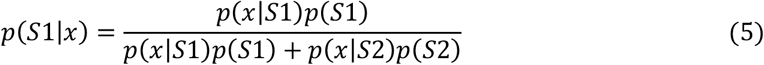

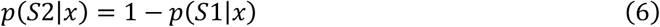

For each trial, objective decision is rendered according to the posterior probability of the S1 state, and confidence is calculated as the posterior probability of the decision being correct.

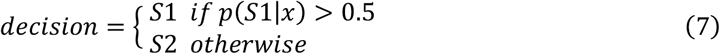

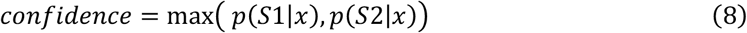

Then, to manipulate the metacognitive performance in simulation data, we have replaced confidence in certain proportions of trials with random samples from a uniform distribution ranging from 0.5 to 1 respectively for S1 and S2 responses (data points in **Figure 7a**). Intuitively, this confidence replacement can be interpreted as the observer completely losing metacognitive information at a certain rate and making a pure guess for confidence rating (i.e., metacognitive lapse rate). The replacement compromises confidence data’s diagnosticity of decision correctness, which appears as an inverted U shape in type-1 zROC space (data points in **Figure 7b**). The simulation data are well captured by the GGSDT (solid curves in **Figures 7a, b**), whose β parameter reasonably reflects the metacognitive disruption yet α_2_ remains constant against the confidence replacement (see **Table 1**, in which σ_1_ and σ_2_ indicate the SD of the S1 and S2 distributions, calculated from α_2_ and β estimates).

**Table 1.** GGSDT parameters estimated in the equal variance simulation.

| replacement | $\mu_1$ | $\mu_2$ | $\alpha_1$ | $\alpha_2$ | $\beta$ | $\sigma_1$ | $\sigma_2$ |
| --- | --- | --- | --- | --- | --- | --- | --- |
| 0% | 0 | 1.052 | 1 | 0.999 | 2.015 | 0.704 | 0.704 |
| 25% | 0 | 1.086 | 1 | 0.994 | 1.601 | 0.815 | 0.810 |
| 50% | 0 | 1.159 | 1 | 1.004 | 1.306 | 0.983 | 0.986 |
| 75% | 0 | 1.309 | 1 | 0.999 | 1.052 | 1.303 | 1.301 |

**Figure 7.**
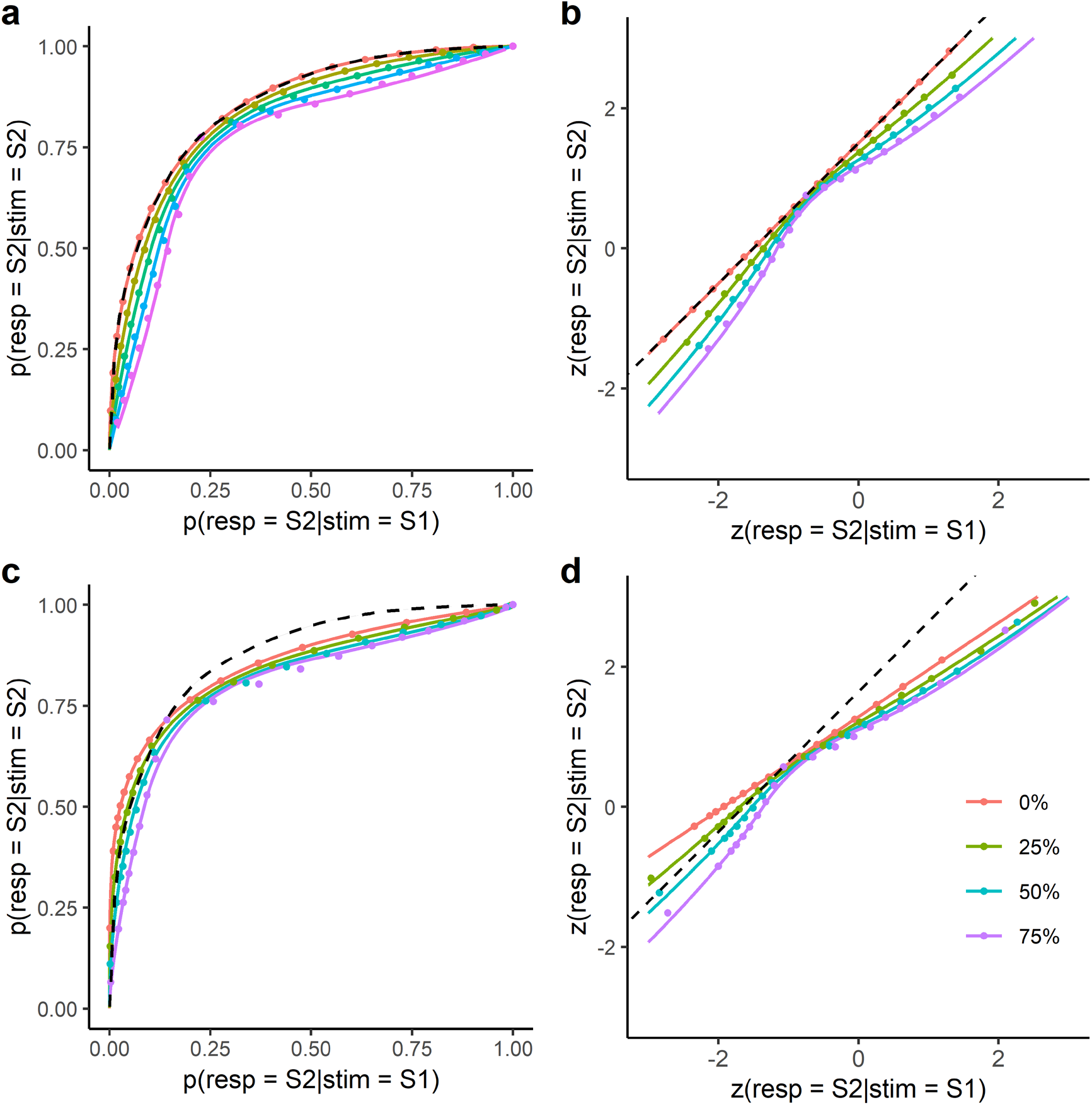
Type-1 ROCs of the GGSDT (solid curves) fitted to simulated data (dots). Data are sampled from the classic equal variance gaussian SDT (**a, b**) or the classic unequal variance gaussian SDT (**c, d**). To implement metacognitive disruption, trial-by-trial confidence data are replaced in 25% steps with samples from a uniform distribution ranging from 0.5 to 1. Data points are plotted using confidence criteria ranging from 0.5 to 0.95 in the step of 0.05. Dashed curves show the prediction of the gaussian meta-SDT of m-ratio = 1.

We also fit the GGSDT to the data sampled from the classic unequal variance gaussian SDT. The internal distributions’ mean distance was set at 1.912 with the SD ratio set to 1.5, which gives the da value of 1.5.^2^ We have sampled 200,000 observations from S1 and S2 distributions and objective decision and confidence rating were rendered by the optimal Bayesian inference. Again, we have replaced certain proportions of confidence with uniform distribution samples ranging from 0.5 to 1 respectively for S1 and S2 responses. The simulation produced asymmetric type-1 ROCs, and the confidence replacement gave the inverted U-shape to zROC (data points **in Figures 7c, d**). The GGSDT nicely captures these data trends (solid curves in **Figures 7c, d**) with reasonable parameter estimates (**Table 2**).

**Table 2.** GGSDT parameters estimated in the unequal variance simulation.

| replacement | $\mu_1$ | $\mu_2$ | $\alpha_1$ | $\alpha_2$ | $\beta$ | $\sigma_1$ | $\sigma_2$ |
| --- | --- | --- | --- | --- | --- | --- | --- |
| 0% | 0 | 1.357 | 1 | 1.508 | 2.006 | 0.706 | 1.064 |
| 25% | 0 | 1.370 | 1 | 1.466 | 1.612 | 0.811 | 1.188 |
| 50% | 0 | 1.409 | 1 | 1.400 | 1.349 | 0.950 | 1.330 |
| 75% | 0 | 1.554 | 1 | 1.333 | 1.094 | 1.230 | 1.640 |

These figures highlight the important difference between the GGSDT and the meta-SDT. If applied to the unequal variance simulation data in **Figure 7c**, the meta-SDT gives considerable misfit due to the equivariance assumption. The dashed curve shows the prediction of the meta-SDT of m-ratio = 1, with the 0% replacement data being above this curve for the S2 responses (points 1st-10th from the left) but below this curve for the S1 responses (points 1st-10th from the right). That is, under the presence of unequal variance, the meta-SDT overestimates (or underestimates) metacognitive efficiency for S1 (or S2) responses.^3^ On the contrary, the GGSDT is not troubled with the unequal variance issue since the heteroscedasticity is captured by the α_2_ parameter while metacognitive efficiency is measured by the kurtosis parameter β, which is assumed to be common for S1 and S2 distributions.

### Interpretability of the β parameter

We have conducted additional analyses to establish the interpretability of β in greater detail. Here, we have simulated classic gaussian SDT data with changing mean distances (1.0, 1.5, 2.0) and SD ratios (1.0, 1.5). For each condition, we have sampled 200,000 observations respectively from S1 and S2 distributions and rendered objective decision and confidence according to the optimal Bayesian inference. To manipulate metacognitive performance, for certain percentages of trials (25% steps), we have replaced confidence with samples from a uniform distribution ranging from 0.5 to 1.

**Figure 8a** plots β values estimated from the simulation data. Strikingly, regardless of the sampling distributions’ mean and SD parameters, β shows a linearly decreasing trend within the confidence replacement rate of 0-0.75. ^4^ Accordingly, within a reasonable parameter range (roughly β > 1), β can be interpreted as a proxy of the metacognitive lapse rate largely irrespective of the internal distribution structure. In other words, β can be understood in reference to the proportion of the gaussian ideal observer’s confidence being disrupted with random guessing. We see the metacognitive lapse rate as an auxiliary construct for a descriptive purpose, yet it may proffer an intuitive characterization of observed metacognitive loss, reinforcing the interpretability of the GGSDT analysis together with the fact that β = 2 represents the same level of metacognitive efficiency as the classic gaussian SDT.

**Figure 8.**
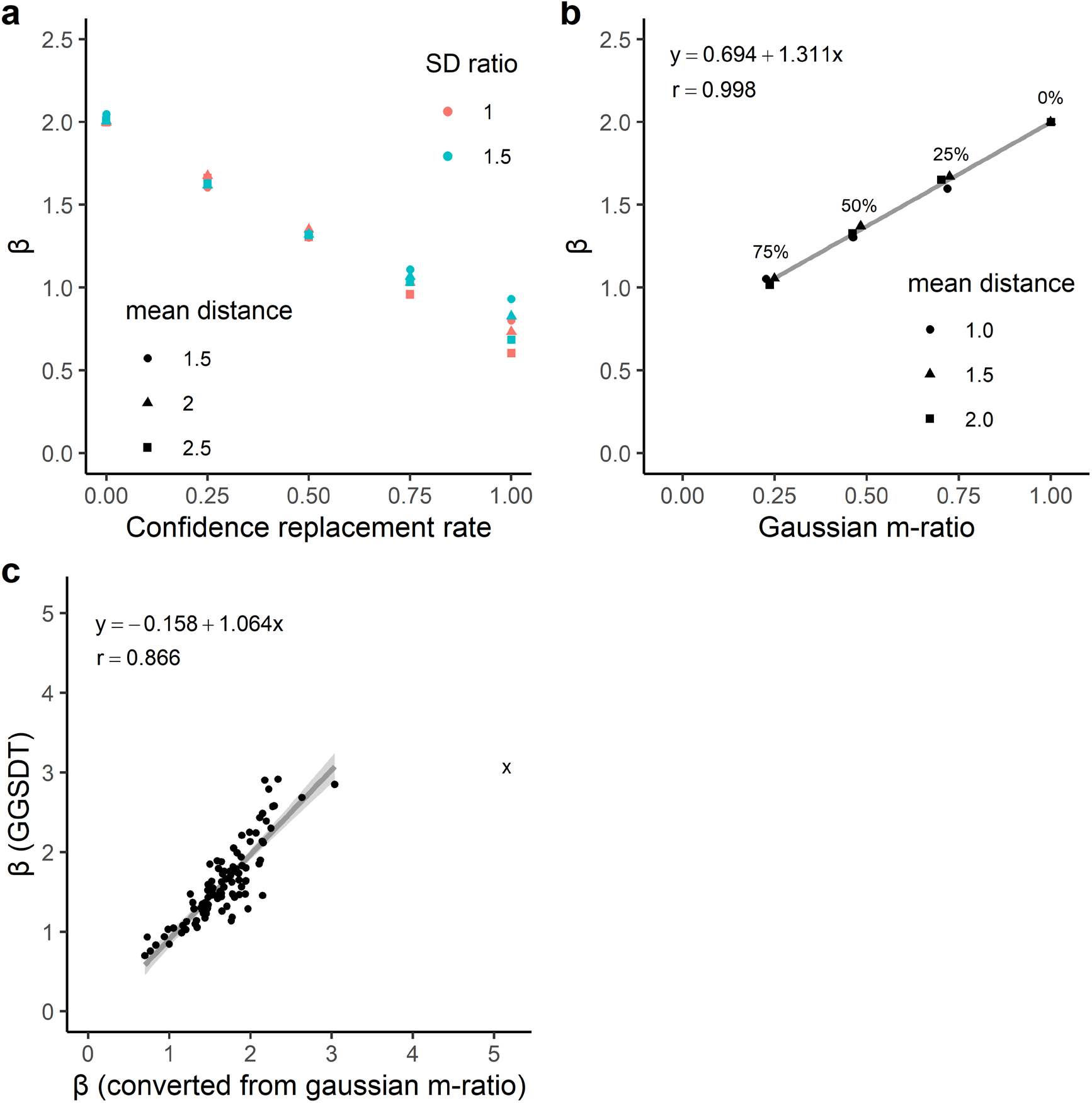
(**a**) Estimated β values under different simulation conditions. Simulation data are sampled from the classic gaussian SDT with varying mean distance (1.0, 1.5, 2.0) and SD ratio (1.0, 1.5), and confidence values are replaced with uniform distribution samples in 25% steps. (**b**) The linkage between the GGSDT’s β and the gaussian meta-SDT’s m-ratio under the equivariance simulation. Simulation data are sampled from the classic equal variance gaussian SDT of different d’ values (1.0, 1.5, 2.0), and confidence was replaced in 25% steps. (**c**) β values estimated for 104 empirical datasets. A linear regression was made after removing one outlier observation marked with the x sign.

Also, under the equivariance situation, the β parameter can be characterized in relation to the meta-SDT framework. **Figure 8b** shows parameter estimates of the GGSDT and gaussian meta-SDT fitted to simulation data sampled from the classic equal variance gaussian SDT. Here, d’ of the sampling distributions was set at 1.0, 1.5, or 2.0, and, respectively for S1 and S2 responses, confidence in certain proportions of trials was replaced with samples from a uniform distribution ranging from 0.5 to 1. Within the confidence replacement rate of 0-0.75, β is almost perfectly correlated with gaussian m-ratio (r = .998), which makes them interchangeable in the equivariance situation by the following formula (see **Appendix 3** for further consideration on this point).

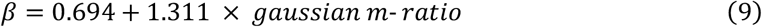

This provides a strong rationale for treating β as a metacognitive efficiency measure being controlled for objective decision accuracy. The intuition behind this is that metacognitive lapse (putative operation of β) causes greater information loss in the situation of higher objective accuracy (when S1 and S2 distributions are less overlapped), through which the influence of objective accuracy is implicitly taken into account in the β estimate (for more discussion about objective accuracy contamination in metacognitive measurement, see Fleming & Lau, 2014; Guggenmos, 2022).

Furthermore, we fit the GGSDT and gaussian meta-SDT to 105 datasets of human 2AFC experiments that were taken from a confidence database (Rahnev et al., 2020) and formerly analyzed in Miyoshi et al. (2022) (see Supplementary Material 1 for details). For each dataset, we have aggregated data from individual participants for estimation stability and estimated β using the GGSDT or the m-ratio conversion with equation 9. The GGSDT model converged for 104 cases, and after removing one outlier observation (marked with x sign in **Figure 8c**), these two variations of β showed a good linear relationship with a slope nearly equal to 1 (r = .866). Therefore, within a reasonable parameter range (roughly β < 4), the crosstalk between these two frameworks holds even in empirical 2AFC experiments, for which the equivariance assumption is not strictly satisfied, thereby ensuring the practical applicability of the GGSDT analysis.

## Application to empirical datasets

### Visual detection versus discrimination

As a next step, we have fitted the GGSDT to empirical datasets selected from the confidence database (Rahnev et al., 2020). We first targeted data from Mazor et al. (2020), in which participants engaged in both yes/no visual detection (asymmetric ROCs are expected) and left/right visual discrimination (symmetric ROCs are expected) for the same set of visual grating stimuli. The GGSDT framework allows direct model-based comparisons of the metacognitive performances between the different formats of decisions.

The dataset includes 46 participants, each engaged in visual detection (presence/absence decision for visual grating) and discrimination (left/right orientation discrimination for visual grating). Each task comprises a total of 200 trials, in which a 6-level confidence rating follows an objective decision. Since the number of available trials is relatively small for each participant, for better model convergence, we have reorganized confidence data into 3 levels by concatenating the confidence rating of 1-2, 3-4, and 5-6. We found 39 cases for which the model converged for both visual detection and discrimination. Further, we have excluded the cases of extreme parameter estimates (α_2_ > 3, β > 4), which have left data from 31 participants for the following analysis (note that the original paper also included only 35 participants for analysis after data exclusion).

**Table 3** shows the averaged parameter estimates. As expected, α_2_ was significantly larger for detection than discrimination, reflecting type-1 ROC asymmetry typically seen for visual detection (t(30) = 4.64, p < .001, **Figure 9a**). There was no significant difference in β between the tasks (t(30) = 0.90, p = .374, **Figure 9b**), and β was not significantly different from 2 in both tasks (t(30) = 0.59, p = .554 for detection, t(30) = −0.76, p = .456 for discrimination). Namely, the participants’ metacognition was not significantly different from the classic gaussian SDT’s prediction, regardless of the decision-making formats.

**Table 3.** GGSDT parameter estimates averaged across 31 participants.

| | $\mu$ | $\alpha_2$ | $\beta$ | $\sigma_1$ | $\sigma_2$ |
| --- | --- | --- | --- | --- | --- |
| Detection | 1.4 (0.18) | 1.47 (0.09) | 2.09 (0.15) | 1.24 (0.3) | 1.78 (0.49) |
| Discrimination | 1.13 (0.07) | 1 (0.04) | 1.9 (0.12) | 0.93 (0.1) | 0.92 (0.1) |

Standard error of the mean is shown in parantheses.

**Figure 9.**
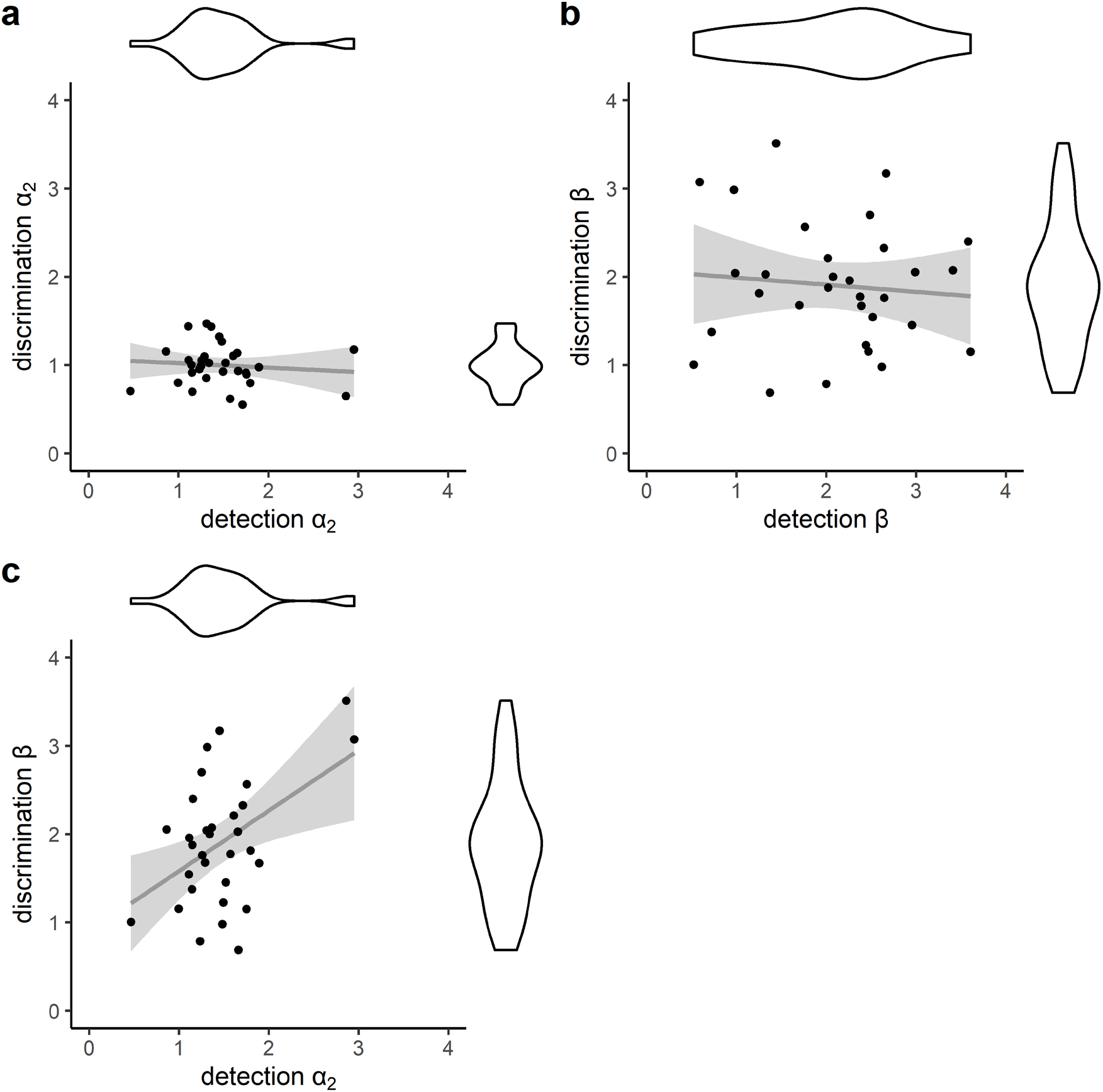
Across-participant correlations of the GGSDT parameter estimates between visual detection and discrimination tasks.

There was no significant across-participant correlation of β between the tasks (r = −.10, t(29) = −0.55, p = .610, **Figure 9b**). This is consistent with previous reports that individual metacognitive performance correlates only within the same task formats but not between yes/no versus 2AFC experiments (Baird et al., 2013; Lee et al., 2018; McCurdy et al., 2013). Interestingly, however, we found a positive correlation between detection α_2_ and discrimination β (r = .471, t(29) = 2.87, p = .008, **Figure 9c**). Of theoretical importance is that this is exactly the pattern predicted by a bidimensional signal detection model of Miyoshi and Lau (2020). More specifically, confidence in the discrimination task is calculated through a heuristic that places a stronger weight on the decision-congruent evidence than the decision-incongruent evidence, the efficiency of which (i.e., discrimination β) is boosted by the unequal variance between target versus nontarget evidence distributions (i.e., detection α_2_). Therefore, the cross-task GGSDT analysis confirms the theoretical perspective of Miyoshi and Lau (2020) and provides a mechanistic insight as to why metacognitive performance hardly generalizes over the different decision formats.

### Analysis of yes/no recognition memory data

Historically, recognition memory research has been the field that uses the zROC analysis most frequently (e.g., Kellen & Klauer, 2018; Osth et al., 2015; Ratcliff & Starns, 2013; Rotello, 2017; Wixted, 2007, 2020). The issue of metacognitive efficiency, however, has not attracted great attention in this field, most likely because studies on recognition memory have usually employed the yes/no design over the 2AFC design, which prevents the application of the meta-SDT analysis. Our use of the GGSDT, which is unrestricted by the equivariance assumption, has allowed us to offer fresh insight into the matter of mnemonic confidence.

We have selected yes/no recognition memory datasets from the confidence database (Rahnev et al., 2020) according to the following criteria: (1) datasets featured with more than 3 levels of confidence rating, (2) datasets whose trial number is no less than 16 x the number of possible response outcomes; e.g., binary decision (hit, miss, false alarm, correct rejection) with 3 levels of confidence gives 12 possible response outcomes. The selection procedure picked up 7 datasets, which include 12 different experimental conditions composed of 653 cases at the individual participant level (see Supplementary Material 2 for details). The GGSDT fitting converged for 598 individual cases, and the removal of extreme parameter estimates (α_2_ > 3, β > 4) has left 444 cases for the following analyses.

**Figure 10a** and **Table 4** summarize the GGSDT parameter estimates. The α_2_ estimate was significantly larger than 1, indicating the heteroscedasticity typically observed in yes/no recognition memory studies (t(443) = 26.82, p < .001). Interestingly, the β estimate was significantly greater than 2, indicating that participants on average were more metacognitively efficient than the classic gaussian SDT (t(443) = 32.78, p < .001). This may suggest that recognition memory behavior can be better modeled with different distributions than the gaussian distribution, which has traditionally been the primary choice in this field (e.g., Miyoshi et al., 2018; Snodgrass & Corwin, 1988; Wixted, 2020). Also, the present recognition memory results contrast with previous findings that gaussian m-ratio < 1 has often been observed in visual perception tasks (e.g., Shekhar & Rahnev, 2020, 2021), suggesting inter-domain idiosyncrasy of human metacognitive efficiency.

**Table 4.** GGSDT parameter estimates averaged across 444 participants.

| $\mu$ | $\alpha_2$ | $\beta$ | $\sigma_1$ | $\sigma_2$ |
| --- | --- | --- | --- | --- |
| 1.08 (0.06) | 1.45 (0.02) | 2.3 (0.04) | 1.37 (0.29) | 1.8 (0.34) |

Standard error of the mean is shown in parantheses.

**Figure 10.**
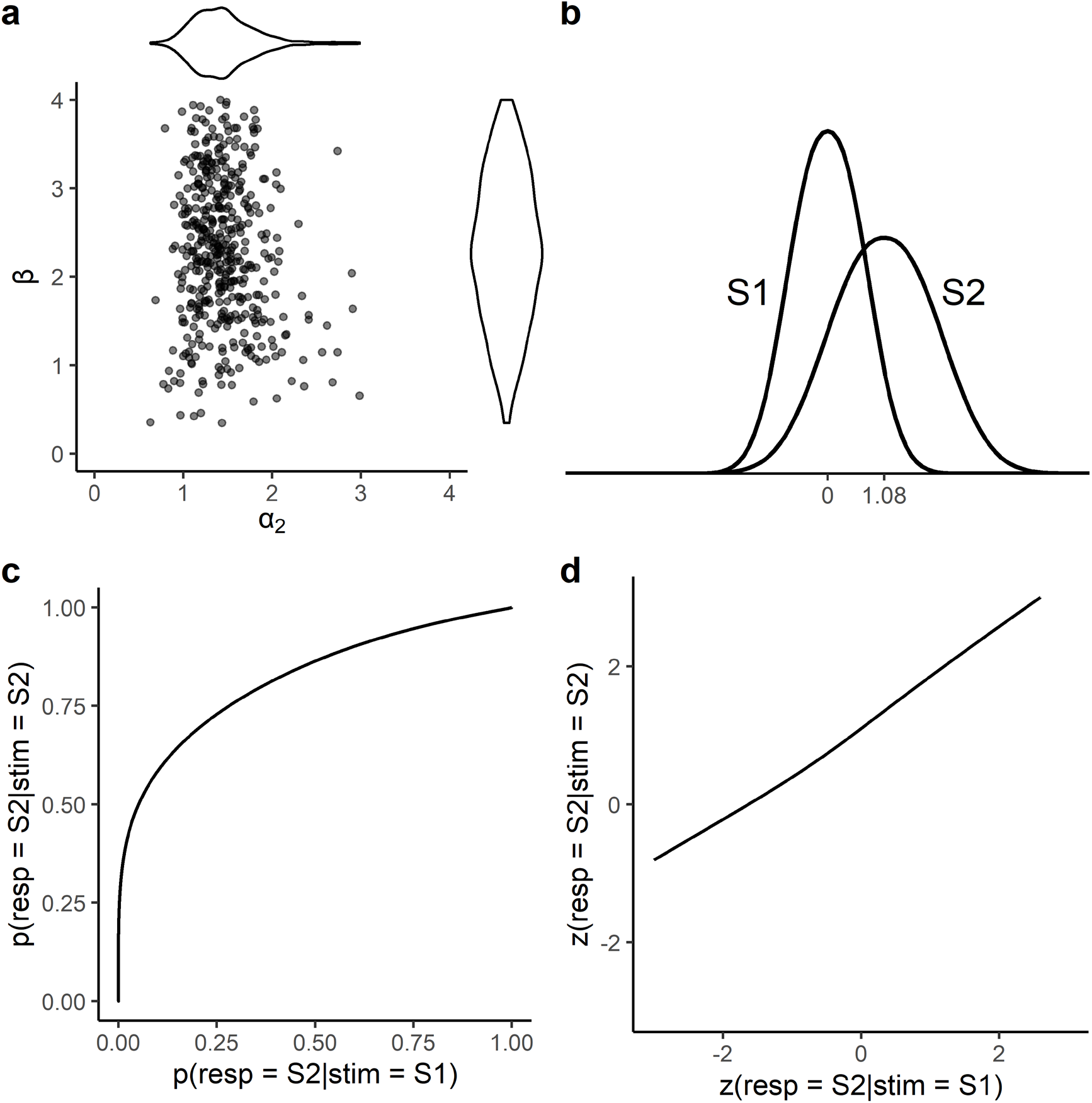
Individual parameter estimates (**a**), and the GGSDT’s behavioral characteristics constructed with the averaged parameter estimates in Table 4 (**b, c, d**).

**Figures 10b-d** show the GGSDT’s decision space and operating characteristics constructed from the averaged parameter estimates in **Table 4**. The internal distributions are featured with relatively small kurtosis (**Figure 10b**), which leads to rather conspicuous patterns of ROCs in **Figures 10c, d**. The type-1 ROC shows a near-vertical rise around the left-most region (almost no false alarms with the highest level of confidence), and the zROC demonstrates U-shaped curvilinearity. Of note is that these patterns closely remind the ROC predictions of the dual-process signal detection model, a hybrid of the discrete threshold process and the continuous signal detection process (Yonelinas, 1997; Yonelinas & Parks, 2007). That is, our GGSDT analysis prescribes the novel evaluation method for zROC curvilinearity (i.e., estimation of β), suggesting that traditional discussions on recognition memory processes could be revisited in light of metacognitive efficiency.

## Discussion

This paper proposes the GGSDT as a novel signal detection framework for analyzing confidence rating data. The GGSDT can evaluate metacognitive efficiency like the meta-SDT, while inheriting the modeling flexibility of the classic SDT to accommodate asymmetric ROCs. The GGSDT quantifies the internal SD ratio and the efficiency of metacognitive judgment based on well-defined statistics (the scale parameter α and the shape parameter β of the generalized gaussian distribution). This unique property would be of great service to empirical performance evaluations as well as artificial data simulations.

As seen in **Figure 8**, β can practically be understood in relation to the proportion of the gaussian ideal observer’s confidence disrupted with random guessing (metacognitive lapse rate), where β = 2 represents the same level of metacognitive efficiency as the classic gaussian SDT. We deem the metacognitive lapse rate to be an auxiliary concept, intended to provide a descriptive data interpretation. In other words, the GGSDT is supposed to be a measurement model, being agnostic of specific processes behind varying levels of metacognitive performance. To pinpoint and draw specific conclusions regarding the mechanisms of particular metacognitive behavior, a process model approach (e.g., Fleming & Daw, 2017; Guggenmos, 2022; Shekhar & Rahnev, 2021; Webb et al., 2022) would be required on top of the current measurement framework.

Our simulations revealed that changing values of β produce upright or inverted U-shaped zROCs, which have been frequently observed in empirical experiments on different cognitive domains (Miyoshi et al., 2022; Shekhar & Rahnev, 2021; Yonelinas, 1997; Yonelinas & Parks, 2007). Therefore, the GGSDT offers a model-based metacognitive measurement while qualitatively conforming with human behavioral characteristics (**Figure 7**). This constitutes a strength of the current framework because inaccurate data descriptions in model fittings may lead to undesirable bias in metacognitive measurement (Dayan, 2022). The GGSDT describes objective decision and confidence by a common set of internal distributions. This, however, does not necessarily imply that they are based on a single decision system. The GGSDT only serves for a posteriori performance evaluation and does not concern itself with whether objective decision and confidence are produced by different processes. As a result, the researcher is free to make whatever presumptions regarding the background mechanisms of the observer’s decision-making behavior. For example, one could assume that decision variables from objective and metacognitive subsystems are combined to constitute unidimensional GGSDT evidence distributions (a similar idea can be seen in a dual-process recognition memory model in Wixted & Mickes, 2010). Alternatively, one can posit that human decision-making behavior is based on multi-dimensional decision variables, and the GGSDT only provides a posteriori unidimensional description of ROC data.

One limitation is that the current GGSDT model cannot accommodate itself to the polygonal type-1 ROC of zero metacognitive efficiency (**Figure 4**) and thus β’s interpretability breaks down under complete metacognitive failure (**Figure 8**). Accordingly, the GGSDT does not completely replace the existing framework of meta-SDT analysis (see **Appendix 3** for more discussion on the compatibility of these frameworks). One reassurance for this is that near-zero metacognitive efficiency is hardly likely to be exhibited by normal human observers (Knotts et al., 2018; Peters & Lau, 2015; Rajananda et al., 2020), and even a blindsight patient GY was reported to show metacognitive performance reasonably above chance (Persaud et al., 2011). Thus, except for the arguably rare scenario of severe metacognitive deficit, the GGSDT’s β offers a reasonable measurement of metacognitive efficiency.

### Ideas and speculation

We propose the GGSDT as a model for measurement purposes. The concept of the metacognitive lapse, however, could be of interest in the future development of sophisticated process models of metacognition. Different sources of metacognitive inefficiency have been considered so far (for a review, see Shekha & Rahnev, 2020) and the metacognitive lapse may be an interesting addition to those. One speculation is that the metacognitive system monitors the activity of multiple neural clusters, many of which are involved in the execution of objective decisions, but some are concerned with unrelated functions. A metacognitive lapse occurs at a certain rate assuming that the confidence level is primarily determined by the input from one of these clusters (e.g., winner takes all of the most active cluster). The idea of the metacognitive lapse, empirically supported by the zROC curvilinearity, may open one intriguing avenue for future research.

## Conclusion

The GGSDT provides a novel measurement method of metacognitive performance that is largely invariant to ROC asymmetry or contamination from objective decision accuracy. Accordingly, this framework can serve a variety of research purposes, including the direct comparisons of visual detection versus discrimination (**Figure 9**) and metacognitive evaluations for asymmetric yes/no recognition memory ROCs (**Figure 10**). The GGSDT analysis can easily be implemented with the *ggsdt* package available from https://github.com/kiyomiyoshi/ggsdt. With its interpretational ease and wide applicability, we hope the GGSDT will be a valuable addition to your analysis toolbox.

## Acknowledgments

This work was supported by JSPS KAKENHI Grant Number 22K13870.

## Materials and methods

### Classic SDT framework

The classic SDT was formalized in the 1950s and has been widely used until the present day (Egan & Clarke, 1956; Green & Swets, 1966; Macmillan & Creelman, 2005; Swets et al., 1955). Here, we explain this framework in reference to yes/no visual detection with 3 levels of confidence (**Figure 1a**), which results in 6 possible response classes (highest confidence S1 to highest confidence S2). **Figure 2a** shows example data plotted in so-called type-1 ROC space, which summarizes cumulative hit rates (S2 response rates to S2 stimuli) and false alarm rates (S2 response rates to S1 stimuli) calculated with different levels of confidence. In counting hits and false alarms, the data points more on the left include trial subsets in which the observer showed greater favor for S2 response. Namely, the left-most data point counts the S2 responses with the highest confidence, the point second from the left concerns the S2 responses with the highest and second-highest confidence, and, eventually, the right-most point includes all the 6 response classes (Macmillan & Creelman, 2005).

**Figure 2b** portrays the classic SDT with unequal variance gaussian distributions assumed for trial-by-trial variability of internal evidence. The distribution on the left represents internal evidence under the S1 state, while the distribution on the right shows evidence under the S2 state. The internal decision space is demarcated by decision criteria (c1-c5) into 6 regions, which represent 6 classes of possible responses; i.e., highest confidence S1 response in the left-most region all the way to highest confidence S2 response in the right-most region. Accordingly, the classic SDT does not explicitly distinguish between objective decision (i.e., S1/S2 response) and confidence ratings in the sense that they are founded on a common set of internal distributions.

Performance evaluation by the classic SDT is made through the fitting to type-1 ROC data. Here, confidence rating data are served for the estimation of the SD ratio of two internal distributions. Usually, the S1 distribution’s mean and SD are fixed at 0 and 1 respectively, and those for the S2 distribution are estimated. A series of decision criteria (c1-c5) is also estimated, and the distribution density delimited by these criteria provides the estimated probability for giving each possible response outcome (e.g., probability of making a false alarm with the confidence level of 3).

There are tractable linkages between the classic SDT’s internal decision space and type-1 ROC space. Unequal variance of the internal distributions (i.e., SD ratio ≠ 1) appears as the asymmetry of the type-1 ROC regarding the major diagonal (**Figure 2c**). To put it another way, the type-1 ROC’s slope at a certain point is given by the likelihood ratio of internal distributions (ratio of the heights of S1 and S2 distributions) at the corresponding location (see **Appendix** 1 for details). Type-1 ROC is sometimes depicted in z-transformed space, where hit and false alarm rates are transformed with the inverse gaussian cumulative distribution function (**Figure 2d**). The classic gaussian SDT gives a straight line in type-1 zROC space, whose slope is determined by the SD ratio of internal distributions. Because of this property, the classic SDT can accommodate asymmetric type-1 ROC data typically observed in yes/no experiments.

The classic SDT prescribes a parametric measure called da, which is the mean difference divided by the root-mean-square SD of two internal distributions (Macmillan & Creelman, 2005, pp. 59-63). The da measure simply indexes the separability between two internal distributions without conceptually differentiating between objective versus metacognitive accuracies; i.e., the classic SDT framework is meant to evaluate the extent to which external world states (S1 or S2) are discriminated together by objective decision and confidence rating.

In summary, the advantages of the classic SDT include the monolithic description of objective decision and confidence rating, the solid connection between its internal distributions and ROC space, and the ability to fit asymmetric type-1 ROC data. This framework, however, does not support the assessment of the observer’s metacognitive accuracy.

### Meta-SDT framework

The meta-SDT evaluates the accuracy of metacognitive monitoring separately from objective decision accuracy. The basic idea was described in Clarke et al. (1959), and the framework was formalized in its current form in Maniscalco and Lau (2012). The meta-SDT posits that objective decision and confidence rating are based on different sets of internal distributions (hereafter called type-1 and type-2 distributions). Here, confidence rating is considered a metacognitive estimate regarding the certainty of one’s own decision correctness (e.g., Mamassian, 2016; Michel, 2022; see also Adler & Ma, 2018; Pouget et al., 2016). This constitutes a stark distinction from the classic SDT, which assumes a common set of distributions for objective decision and confidence rating.

We will use left/right 2AFC discrimination with 10 levels of confidence to demonstrate the meta-SDT analysis (**Figure 1b**). **Figure 3a** illustrates example data plotted in type-1 ROC space. The data can also be plotted in so-called type-2 ROC space, which summarizes the type-2 hit rate (high confidence rate for correct objective decisions) and the type-2 false alarm rate (high confidence rate for incorrect objective decisions) with different levels of confidence (**Figure 3b)**. In counting high confidence responses, the left-most data point only refers to the highest level of confidence, the point second from the left concerns the highest and the second-highest confidence, and so forth. Type-2 ROC represents the observer’s metacognitive accuracy since the area under the curve becomes larger as confidence rating becomes more diagnostic of the correctness of objective decision.

Here, the meta-SDT model adopts equal variance gaussian distributions for both type-1 and type-2 distributions (**Figures 3c, d**). This equal variance assumption is essentially mandatory, and thus the meta-SDT analysis is only appropriate for the situation signified with symmetric S1 versus S2 distributions (i.e., data that are symmetric in type-1 ROC space). Such patterns of data are expected in those experiments that offer symmetrical treatment to S1 and S2 stimuli (e.g., genuine 2AFC paradigm, left/right motion discrimination, etc.) than those of asymmetric treatment (e.g., yes/no paradigm, etc.).

The fitting of type-1 distributions is made by only considering objective decision outcomes (a pair of hit and false alarm rates) without taking confidence data into account. Namely, the following formula estimates the standardized distance of type-1 distributions known as d’, where z() stands for the inverse gaussian cumulative distribution function.

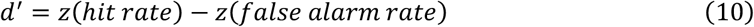

Then, under some parametric constraints, type-2 distributions are fitted to type-2 ROC data. Here, estimated is the standardized distance between type-2 distributions, known as meta-d’ (for technical details, see Barrett et al., 2013; Maniscalco & Lau, 2012). Conventionally, d’ and meta-d’ are directly compared as they are expressed on the same scale (standardized mean difference of equivariance distributions). Accordingly, their ratio (meta-d’/d’, m-ratio) or difference (meta-d’ - d’, m-diff) are often employed to quantify the observer’s type-2 versus type-1 processing efficiency (i.e., metacognitive efficiency, a term for representing metacognitive performance controlled for objective decision accuracy, see Fleming & Lau, 2014).

In short, unlike the classic SDT, the meta-SDT allows for the evaluation of metacognitive efficiency, but its use is restricted by the essential premise of equal variance. This constituted a primary motivation for us to develop the GGSDT as a metacognitive measurement model of broader applicability. Also, the meta-SDT analysis needs to be interpreted with caution because it is sensitive to auxiliary distributional assumptions (Miyoshi et al., 2022). The model of smaller distribution kurtosis (e.g., gaussian meta-SDT) fundamentally gives smaller m-ratio than the model of greater kurtosis (e.g., logistic meta-SDT). It is this kurtosis dependency that gave us a mechanistic insight to design the GGSDT analysis.

### Appendix 1

#### How metacognitive accuracy is mapped onto type-1 ROC

Let us consider plotting type-1 ROC from decision variable distributions provided under S1 and S2 states (**Figure A1**). What is important here is that the type-1 ROC’s slope at a certain point is determined by the density ratio of S2 versus S1 samples at a corresponding location of the decision variable space.

As a concrete example, **Figure A1a** shows the histogram and density curve of the decision variable x, sampled from the equal variance gaussian SDT (d’ = 1). The density ratio of the S2 versus S1 samples (**Figure A1b**) shows how the type-1 ROC slope changes according to the value of x (**Figure A1c**). Here, for simplicity, we can consider x = 0 as the objective decision boundary, which gives hit and false alarm rates of (.31, .69), located in the middle of the type-1 ROC curve. We can also consider the x’s absolute distance from 0 as the confidence value. In this way of thought, the type-1 ROC’s expanding curvature seen in **Figure A1c** can be interpreted as a reflection of the confidence-accuracy relationship (i.e., metacognitive accuracy) prescribed by the equal variance gaussian SDT of d’ = 1. In general, when confidence has positive diagnostic power for decision correctness, type-1 ROC data, other than the objective decision data point ([.31, .69] in the above case), demonstrates expanding curvature.

For another example, let us look at a case with the same objective decision performance as above, but where the confidence (absolute value of x) has no diagnostic power at all for decision correctness (**Figure A1d**). This can be seen from the fact that the density ratio of the correct and incorrect samples is always constant, regardless of the value of x. In other words, the density ratio of the S2 versus S1 samples is a stepwise function (**Figure A1e**), which gives a polygonal type-1 ROC segmented at (.31, .69) (**Figure A1f**). Namely, this type of polygonal ROC is indicative of complete metacognitive failure (besides, this could inspire a nonparametric metacognitive efficiency measure such as the observed type-1 ROC area contrasted with the polygonal ROC area of zero metacognitive accuracy).

Taking it one step further, we see that representing complete metacognitive failure requires irregularly shaped distributions like the ones in **Figure A1d** (to be precise, any distributions giving a constant correct versus incorrect density ratio serve this purpose). Assuming that the decision variable distributions employed by actual human observers are not very far from symmetric bell-shaped distributions, it makes sense that complete metacognitive failure has rarely been observed in empirical experiments, except when d’ itself is very small; even a blindsight patient is reported to have non-zero metacognitive accuracy (Persaud et al., 2011). This line of thinking gives some credit to the GGSDT framework, which aims to evaluate metacognitive performance by the kurtosis of the symmetric bell-shaped distributions.

**Figure A1.**
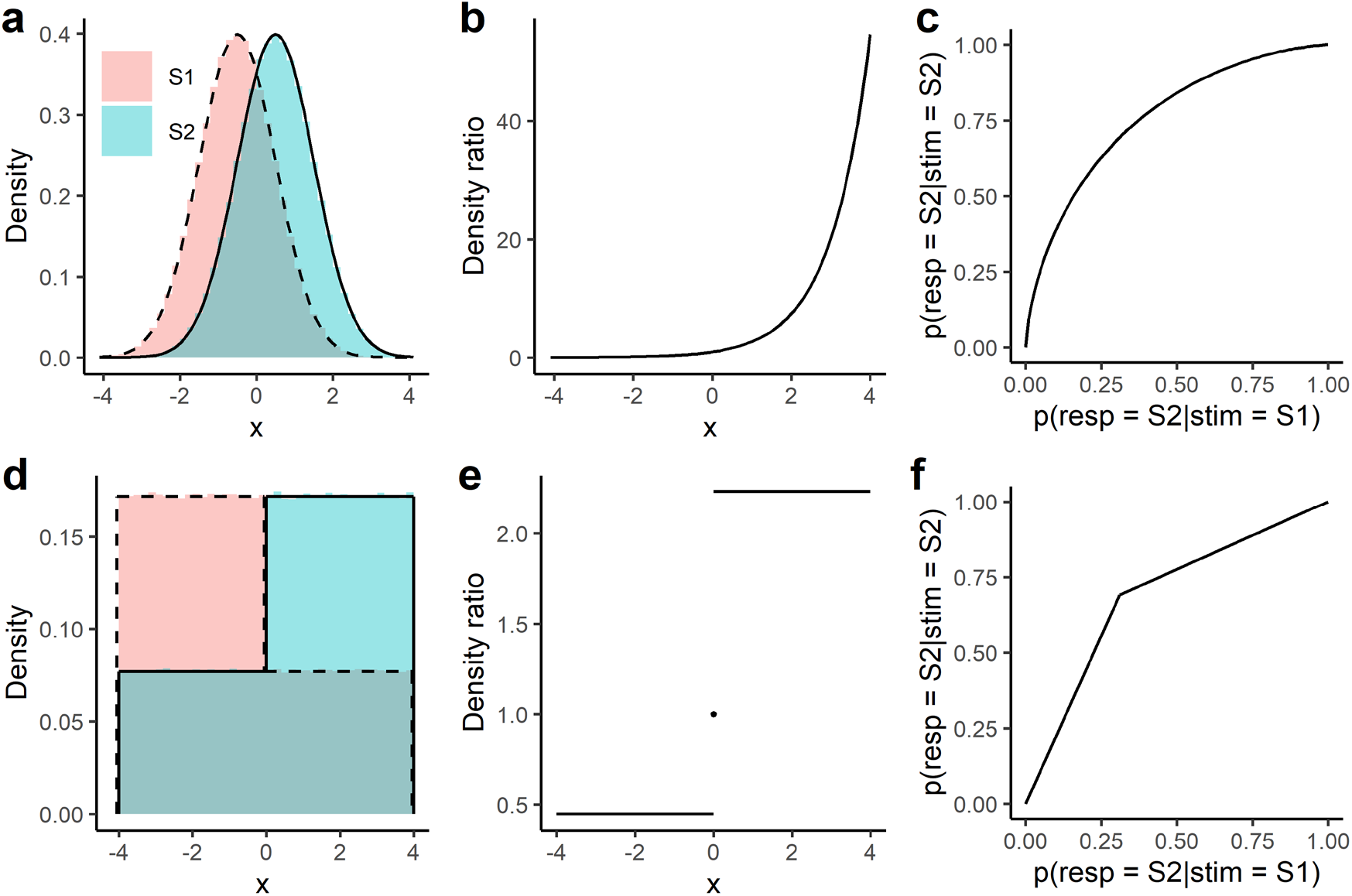
Transformation of decision variable distributions to type-1 ROC. Histograms and density curves of decision variable samples (**a, d**). Density ratio of S2 versus S1 samples (**b, e**). Type-1 ROC constructed from the S1 and S2 density curves (**c, f**).

### Appendix 2

#### Behavioral characteristics of the GGSDT

**Figures A2 and A3** demonstrate the GGSDT’s behavior under varying β values (α_2_ is set at 1 and 1.5 respectively). For both figures, μ_2_ was determined so that the standardized mean difference (da) would be 1.5 under the β value of 2.0 (i.e., baseline condition illustrating the classic gaussian SDT’s behavior). The GGSDT shows different type-1 characteristics according to the α_2_ parameter. However, regardless of the α_2_ values, type-2 characteristics are arguably constant under a given value of β. This constitutes another example of β’s SD-ratio-independency. This property may particularly be useful when one generates simulation data to manipulate both metacognitive efficiency and ROC asymmetry.

**Figure A2.**
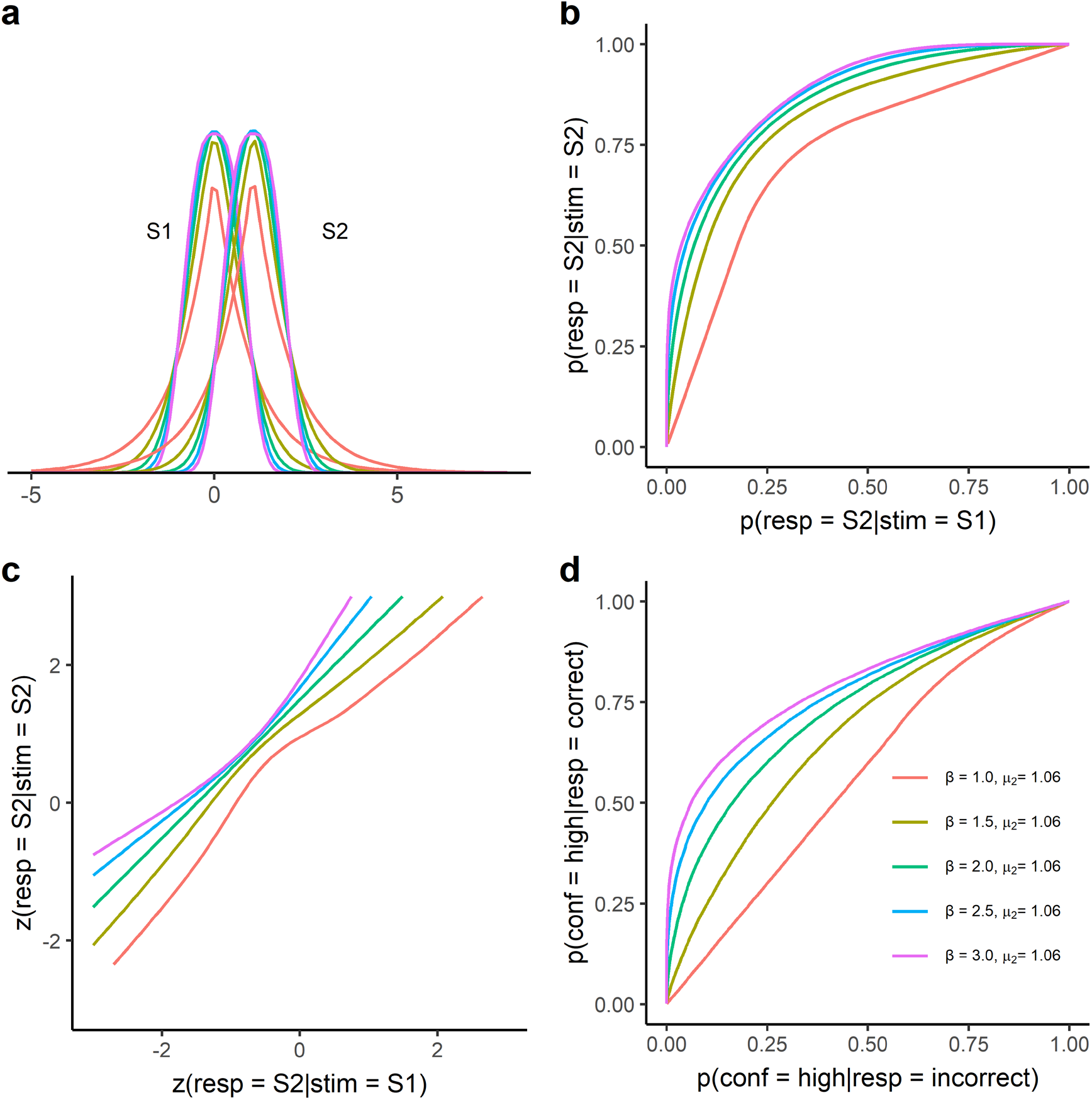
The behavior of the GGSDT under the equal variance scheme (α_2_ is set at 1.0). (**a**) Internal decision space. (**b**) Type-1 ROCs. (**c**) Type-1 zROCs. (**d**) Type-2 ROCs.

**Figure A3.**
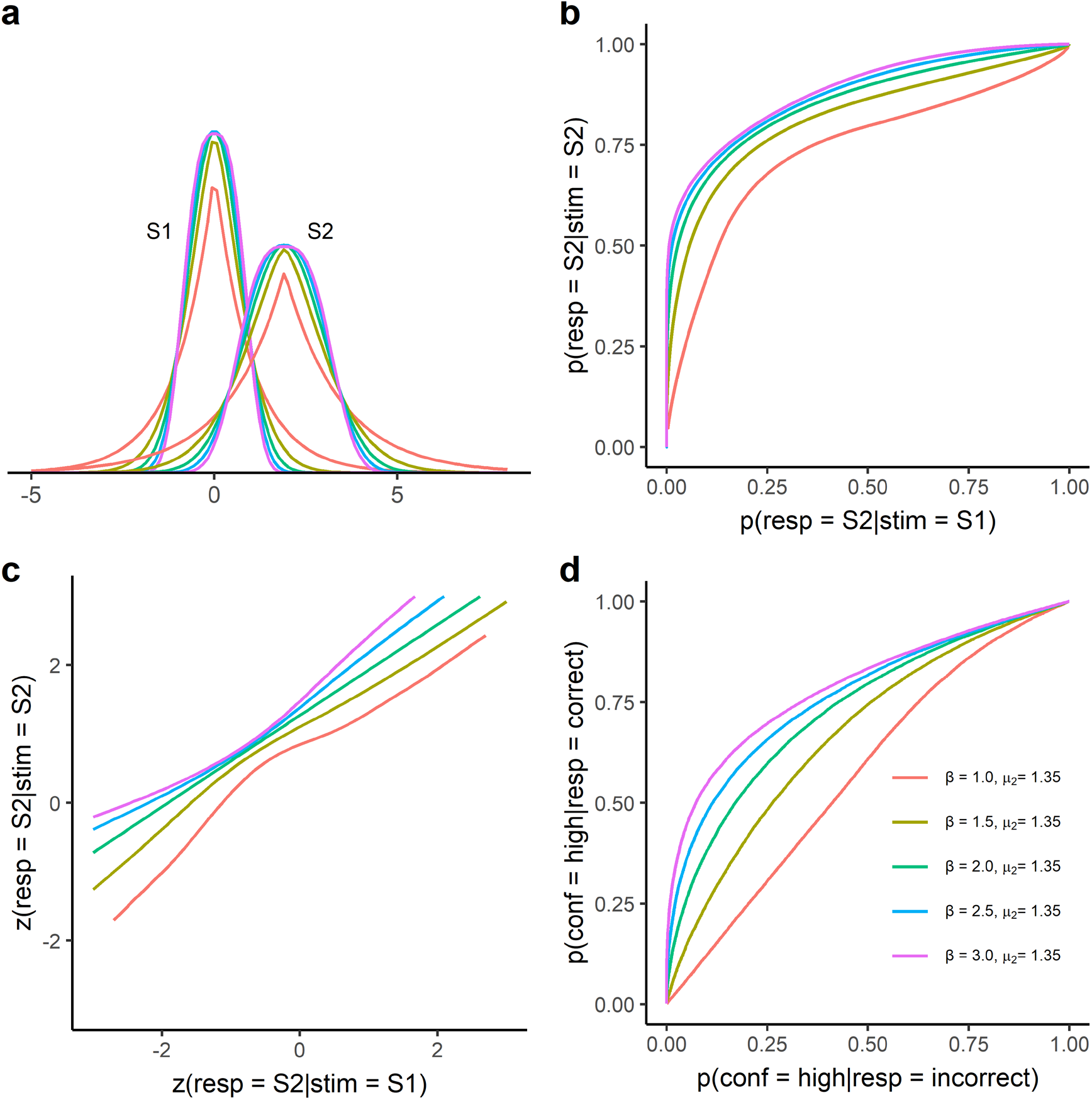
The behavior of the GGSDT under the unequal variance scheme (α_2_ is set at 1.5). (**a**) Internal decision space. (**b**) Type-1 ROCs. (**c**) Type-1 zROCs. (**d**) Type-2 ROCs.

### Appendix 3

#### The crosstalk between GGSDT and meta-SDT

**Figures 7 and 8** show the compatibility of the GGSDT and meta-SDT frameworks based on the simulation data of random confidence replacement. Strictly speaking, however, this metacognitive loss of random guessing (metacognitive lapse) is not perfectly consistent with the measurement regime assumed by the meta-SDT model. The meta-SDT model is supposed to be a measurement model and is widely deployed for practical use even if data structure does not perfectly fit its modeling assumptions (the same can be said for the GGSDT). Still, it may be helpful to fit the GGSDT to the data from the meta-SDT regime further to examine the compatibility of the β and m-ratio measures.

The points in **Figures A4a, b** display simulation data of varying gaussian m-ratios generated in a way that is consistent with (and thus perfectly explained by) the gaussian meta-SDT model (d’ was fixed at 1.6). Namely, we have first simulated objective decisions according to the equal variance gaussian SDT of d’ = 1.6 (see equations 5-7), which gives a middle objective data point in type-1 ROC space. Then, maintaining the obtained number of hits and false alarms, we have simulated the other type1 ROC data points under the equal variance gaussian SDT of different d’ values (0.4, 0.8, 1.6, and 2.0) according to equation 8. The solid lines demonstrate the fitting curves of the GGSDT, which provide reasonable fits to the meta-SDT data (see **Figure 7** for a comparison). One interesting observation is that the meta-SDT data exbibit the U-shaped curvilinearity in the type-1 zROC space, as with the case of the GGSDT. This indicates that the meta-SDT and GGSDT prescribe performance evaluations under qualitatively similar measurement regimes.

**Figure A4c** shows the GGSDT’ β parameter estimated from the simulation data. The solid line represents the mutual parameter transformation defined by equation 9. Although this equation is derived in the confidence replacement simulation (see **Figure 8**), it still gives a good explanation for this data set of the meta-SDT regime. It is also of note that equation 9 gives a good parameter transformation even for the case of the gaussian m-ratio > 1 (i.e., β > 2), which could not be demonstrated in **Figure 8**.

In sum, although the GGSDT and meta-SDT have different measurement regimes in a precise sense, there is good consistency in their operating properties (represented by the curvilinear zROC), and they both would be of practical use while complementing each other.

**Figure A4.**
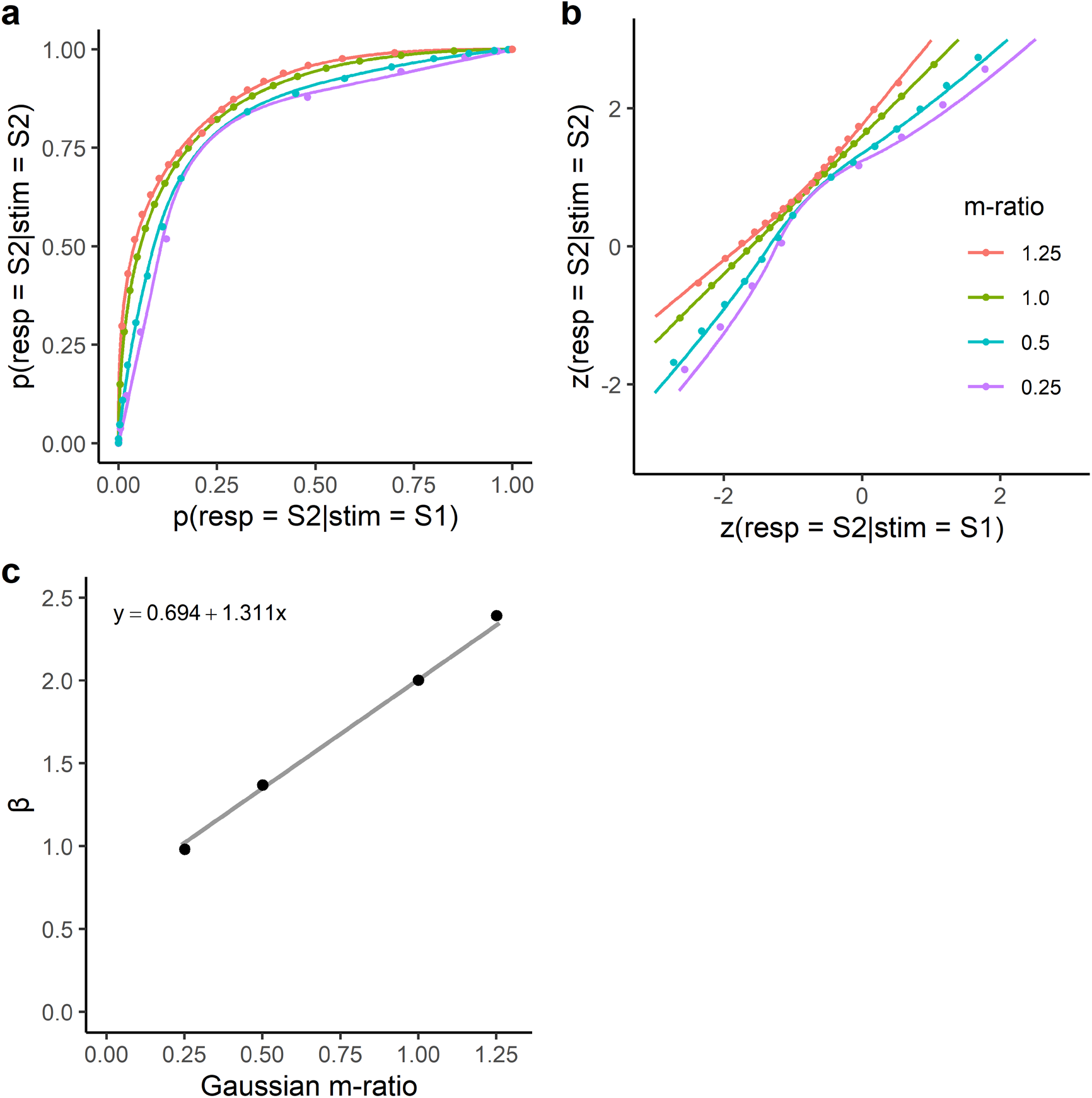
(**a**) Type-1 ROC data simulated with the gaussian meta-SDT regime (data points) and fitting curves of the GGSDT model (solid lines). (**b**) Type-1 zROC data of the gaussian meta-SDT regime (data points) and fitting curves of the GGSDT model (solid lines). The d’ value is fixed at 1.6 for all the cases. (**c**) The GGSDT’s β estimated from the simulation data. The solid line shows the parameter transformation defined by equation 9.

## Footnotes

1 We see β as a measure of metacognitive efficiency since it is shown to be interpretable without much regard to objective decision accuracy (see later sections).

2 The da measure is an extension of d’ for unequal variance and reduces to d’ in the case of equal variance (see **Materials and methods**).

3 To avoid the misfit, researchers sometimes fit the meta-SDT respectively to S1 and S2 responses and estimate response-specific meta-d’ (Maniscalco & Lau, 2014). If this method is applied to the simulation data in **Figure 7c**, meta-d’ is estimated to be larger for S2 than S1 responses, simply reflecting the variance discrepancy between S1 and S2 distributions. This is qualitatively different from the GGSDT analysis, which measures metacognitive efficiency in terms of distribution kurtosis independently from the SD-ratio of S1 and S2 distributions.

4 This linear relationship breaks down by the 100% confidence replacement. This should be due to the limitation that a polygonal type-1 ROC of zero metacognitive efficiency (**Figure 4a**) cannot be represented by a generalized gaussian distribution featured with bell-shaped symmetry. Since near-zero metacognitive efficiency is hardly likely to be observed in empirical experiments, this would not be a major problem in practical use. Note that an irregularly shaped distribution such as seen in the low-threshold model (Kellen et al., 2016; Luce, 1963; Macmillan & Creelman, 2005, pp. 86-88) is necessary for representing the polygonal type-1 ROC of complete metacognitive inefficiency (see **Appendix 1**).

## Notes

### Competing Interest Statement

The authors have declared no competing interest.

### Summary of Updates

Figure 1 has been corrected.

